# The early adolescent brain on music: analysis of functional dynamics reveals engagement of orbitofrontal cortex reward system

**DOI:** 10.1101/2020.06.18.148072

**Authors:** M. C. Fasano, J. Cabral, A. Stevner, P. Vuust, P. Cantou, E. Brattico, M. L. Kringelbach

**Affiliations:** Department of Psychology and Behavioural Sciences, Aarhus University, Aarhus, Denmark; Department of Psychiatry, University of Oxford, Oxford, United Kingdom; Life and Health Sciences Research Institute (ICVS), School of Medicine, University of Minho, Braga, Portugal; Center for Music in the Brain, Aarhus University & The Royal Academy of Music Aarhus/Aalborg, Aarhus, Denmark; Department of Education, Psychology, Communication, University of Bari Aldo Moro, Bari, Italy; Centre for Eudaimonia and Human Flourishing, University of Oxford, UK

**Keywords:** adolescence, music listening, dynamic functional connectivity, orbitofrontal cortex, pleasure

## Abstract

Music listening plays a pivotal role for children and adolescents, yet surprisingly few neuroimaging studies have studied the underlying functional dynamics. We used functional magnetic resonance imaging to scan 17 preadolescents aged 10-11 years old while listening to music. We subsequently tracked the occurrence of functional brain networks over time by using a recent method that detects recurrent BOLD phase-locking states: the Leading Eigenvector Dynamics Analysis (LEiDA). In particular, we compared the probabilities of occurrence and switching profiles of different patterns of BOLD phase-locking between music and no music. Moreover, we used an adapted version of the Barcelona Music Reward Questionnaire (BMRQ) to measure the music reward sensitivity of the participants. Our results showed significantly increased occurrence of a BOLD phase-locking pattern during music listening compared to no music, characterized by a phase-shift in the BOLD signals of the medial orbitofrontal and ventromedial prefrontal cortices – a brain subsystem associated to reward processing – from the rest of the brain. Moreover, we observed a significantly higher probability of switching to this pattern while listening to music. We also found a positive correlation between the individual musical reward sensitivity and the tendency to switch to this reward state during music. Our findings highlight the involvement of a brain subsystem involved in hedonic processing during music listening in the early adolescent brain. These results offer novel insight into the neural underpinnings of musical reward in early adolescence and may help us to understand the importance of music at this delicate age.

## Introduction

Music listening is an important source of enjoyment and entertainment for children and adolescents (Giacometti et al. 1981; North et al. 2000; Erkkila and Saarikallio 2007; Miranda and Claes 2009; Roberts et al. 2009). The increased tendency to listen to music during adolescence coincides with the delicate transition from childhood to adulthood – a critical life phase characterized by developmental and psychosocial challenges and reward-seeking behaviors (Steinberg and Lerner 2004; Miranda and Claes 2009). In particular, while the bottom-up limbic structures including nucleus accumbens and amygdala, implicated in incentive and emotional processing, undergoes dramatic development during adolescence (Laviola et al. 2001; Ernst et al. 2005; Galvan et al. 2006), top-down control systems residing in prefrontal regions matures slowly, generally up to the mid-20 years of age (Gogtay et al. 2004; Kringelbach 2005; Giedd 2008; Asato et al. 2010; Tamnes et al. 2010). Accordingly, adolescence is characterized by heightened sensitivity to rewards paired with minimal cognitive control and emotion regulation (i.e., inability to delay rewards), and thus high impulsivity (Winstanley et al. 2006; Dahl and Gunnar 2009; Chein et al. 2011; Whelan et al. 2012; Fasano et al. 2019). In line with this, pleasure-inducing activities, such as music listening, peak in adolescence (Bonneville-Rouss 2013; North et al., 2000). Some studies showed the importance of music listening in adolescence for developmental tasks and psychosocial adaptive functions, such as self-actualization, individual and cultural identity, socialization and integration with peers, and, above all, emotion regulation (Giacometti et al. 1981; Russell and A. 1997; North et al. 2000; Tarrant et al. 2000; Schwartz and Fouts 2003; Berns et al. 2010). This raises an urgent need for exploring the neural underpinnings of these beneficial effects that music listening is able to produce at this age.

Recently, the *free listening* paradigm has been introduced in fMRI music research which has allowed for the investigation of the neural underpinnings of music listening in a naturalistic situation (Jola et al., 2013; Fasano et al. n.d.; Abrams et al. 2013; Hasson et al. 2004; Alluri et al. 2012; Burunat et al. 2014; Silbert et al. 2014; Toiviainen et al. 2014) highlighting the recruitment of auditory, parietal, frontal, subcortical and motor areas. More recently, the advent of network neuroscience has brought insights into the functional connectivity (FC) between brain regions during the listening of musical pieces (Wilkins et al. 2014; Karmonik et al. 2016; Alluri et al. 2017; Toiviainen et al., 2019), showing that real-world music listening is associated with neural connectivity patterns rather than activity in isolated regions (Pfeifer and Allen 2012; Reybrouck et al. 2018). However, despite the important role that music listening plays in the lives of children and adolescents, none of the previous neuroimaging studies has so far investigated the neural underpinning of naturalistic music listening (not interrupted by behavioral tasks) in this young population.

In the present study we aim at investigating the brain networks involved in naturalistic music listening in early adolescence focusing on the temporal dynamics of a set of recurrent functional networks. While the scarce research that has been conducted on FC during music listening has been focused on ‘static’ FC, representing mean connectivity over a period of scanning, growing evidence shows indeed that brain FC is not stable during the scan, but instead exhibits a repertoire of timevarying, but reoccurring, states of coupling among brain regions (Deco et al. 2011; Hansen et al. 2015; Cabral, Kringelbach, et al. 2017; Vohryzek et al., 2020). Therefore, we used a sensitive neuroimaging analysis method, the Leading Eigenvector Dynamics Analysis (LEiDA), which defines FC based on whole-brain BOLD *phase-locking* patterns (Cabral et al. 2017). Unlike other methods to detect FC dynamics based on BOLD co-activation patterns, which rely mainly on the BOLD amplitude dynamics (Karahanoğlu and Van De Ville 2015), LEiDA exhibits high sensitivity to capture slow BOLD oscillatory modes characterized by both co-activations and co-deactivations. Clustering the patterns of BOLD Phase Locking (PL) allows identifying PL states that reoccur over time across different scanning sessions (Cabral et al. 2017; Vohryzek et al., 2020; Stark et al., 2020; Lord et al., 2019; Larabi et al., 2020). By using LEiDA, we wanted to evaluate if there are specific BOLD PL configurations that differentiate music listening from no music conditions in children entering adolescence. Furthermore, under the hypothesis that these BOLD PL patterns may relate with the hedonic experience of music listening, we correlate their occurrence with the music reward sensitivity of the young participants.

## Methods and Materials

### Participants

Participants were part of a longitudinal study addressing the impact of music training on psychological, pedagogical and neural functions and were recruited from public elementary schools in Aarhus district (Denmark). We recruited 17 preadolescents 10-11 years of age (mean age: 10 years and 7 months; SD: 5,8 months) including ten girls and seven boys, all with normal hearing. Exclusion criteria included any history of developmental or neurologic disorders. The parents of the participants were informed about the possibility to participate in the project with an invitation sent to them via intranet and e-mail by the schools, and via direct communication with the teachers and the project personnel.

### Experimental procedure

Study protocols were approved by the official Midtjyllands Regional Science Ethics Committee (1-10-72-122-16). Written informed consent for participation in the study was obtained from the parents/guardians on behalf of the child participants and verbal assent was obtained from all children individually. Either the parents or the children could end their participation at any time. Parents/guardians received voucher compensations for their child’s participation in the form of cinema tickets and Amazon vouchers and children were awarded small prizes (e.g. toys or stickers). All preadolescents were tested individually at Aarhus University Hospital. The psychological tests used for the longitudinal project in which this study is included where administered at the schools attended by the children, with previous agreement of the headmasters. For the scanning session, we designed a child-friendly protocol that included a 30-minutes preparation prior to the actual scanning. Preadolescents were carefully informed about what a scanner is, what they would do during the whole scanning time, and how important it was to stay still, with the help of some pictures and simulations. They also had some time to explore the scanner room and to test the bed for a brief familiarization period. Before starting the scanning session, all the participants practiced staying still and were exposed with some of the sounds made by the scanner. To avoid any distress for the child, the parents were given the possibility to accompany the child in the MR room and, eventually, remain in the scanner room and held their hand. Preadolescents watched a cartoon movie during the anatomical scan to assist them with staying still.

### Behavioral assessment

The administration of psychological tests and questionnaires had a total duration of 45 minutes. Only the questionnaires relevant for this study are reported and discussed here.

#### BMRQ

We used a Danish version for children and adolescents of the Barcelona Musical Reward Questionnaire (BMRQ). The BMRQ (Mas-Herrero et al., 2013) is a psychometric instrument known to be a reliable indicator of interindividual variability in music-induced reward. Consistently with previous neuroimaging studies (Martínez-Molina et al. 2016), we considered the BMRQ overall scores for our analysis.

#### Music training

We collected information about the individual level of music experience by asking the participants and their parents to indicate the number of months of music training received.

#### Socio-economic status

Parents were asked to indicate their highest level of education and annual household income on a questionnaire. Responses to education level were scored on a 5-point scale: 1) elementary/middle school; 2) high school; 3) college education; 4) master’s degree; 5) professional degree. Responses to annual household income were scored on a five-point scale: 0) < DKK 65399; 1) DKK 65400-130699; 2) DKK 130700–195999; 3) DKK 196000–261399; 4) DKK 261400– 326699 and 5) > DKK 326700. A final socio-economic status (SES) score was calculated as the mean of each parent’s education score and annual income, consistently with previous studies (Habibi et al. 2014; Sachs et al. 2017).

### Stimuli

The stimuli used for the fMRI paradigm were two medleys of violin pieces for children (referred to as piece 1 and piece 2 in the rest of the manuscript), each with a duration of 3 min and 50 s, for a total of 7 min and 40 s. The pieces included in the medleys were: “*Winter time in Russia*” by Joanne Martin; “*Lift off*”, “*Tiptoe boo!*”, “*Here it comes*”, “*In flight*”, “*Clare’s song*” by K. & D. Blackwell, “*E-, A-, D-, G-Stränglåten*” by Eva Bogren. We selected violin pieces for children because this study was the first part of a longitudinal project including two fMRI measurements and looking at the effect of music learning, therefore requiring the music stimuli to be unfamiliar to all participants and easy to be learned over half a year. Moreover, we used two pieces in order to compare, after the learning period, the neural response while listening to a learned (piece 1) vs. not learned (piece 2) piece of music. Here we present the results of the first fMRI measurement when both the pieces were unfamiliar to the participants. Fourteen pieces where listened to, assessed and rated for a list of technical aspects by three professional violinists and, subsequently, 4 couples of similar pieces were chosen by two of the experimenters (MCF and EB) - who are both professional trained musicians - in order to create two medleys (piece 1 and piece 2) matched as much as possible for complexity, tempo, genre and accompaniment. The order of piece 1 and piece 2 was counter-balanced across subjects. Before the presentation of the two stimuli, which is the focus of the current study, the participants were familiarized with the *free listening* task by listening to a brief excerpt of music with a duration of 29.9 s. All the audio stimuli were preceded and followed by 29.9 s of no music for a total of 2 min of no music (Figure 1D, *middle*). At the end of the *free listening* paradigm, the preadolescents where asked how much they liked each of the two pieces of music.

**Figure 1.**
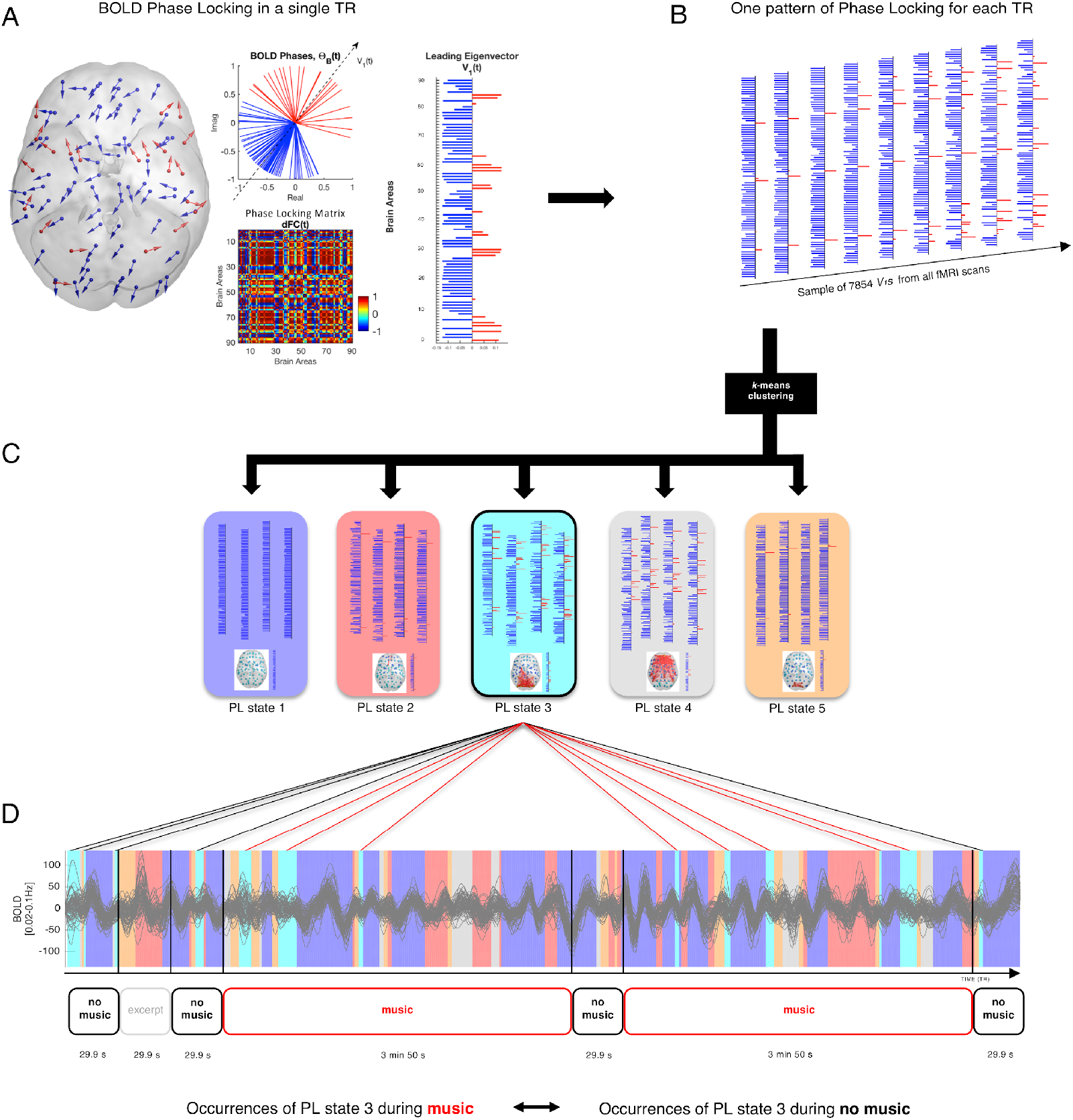
Illustration of the LEiDA methodology to detect recurrent states of BOLD Phase Locking and compare their probability of occurrence between 2 conditions (here music vs no music). **A –** *Left and middle:* at each single time point, the BOLD phases in all N=90 brain areas are represented both in the cortical space (arrows placed at the center of gravity of each brain area) and in the complex plane, where all phases are centered at the same origin. The phaselocking matrix at time *t* captures the BOLD phase-locking between each pair of brain areas, where 1 means full synchrony and −1 indicates a phase difference of 180°. *Right:* the leading (i.e. largest magnitude) eigenvector of the phase-locking matrix at time *t*, *V_1_(t)*, is the vector that best captures the orientation of all BOLD phases, where each element in *V_1_(t)* corresponds to the projection of each area BOLD phase into *V_1_(t)*. The signs of the elements in *V_1_* (red/blue) are used to divide brain areas into communities according to their BOLD phase. **B –** The leading eigenvectors (*V_1_*s) are obtained for each time point from all fMRI scans in all subjects, resulting in a large sample of 7854 leading eigenvectors. **C –** This sample is partitioned into a reduced number of *K* clusters. For illustration, a partition model (clustering solution) of 5 PL states is considered. Each cluster is represented by a central vector. We take these cluster centroid vectors as representing recurrent patterns of BOLD phase locking, or PL states. Each of these centroid vectors is here illustrated as a network in cortical space. Here lighter and darker colors show stronger and weaker coherence, respectively, within the positive (yellow-red) vs. the negative (cyan-blue) communities. **D –** *Top:* to obtain the PL state time courses, we select, at each TR, the centroid to which *V_1_(t)* is the most similar. In this way, for each fMRI session, one PL state is assigned to each TR (highlighted in the same color of the corresponding recurrent state). For illustration, the PL state assigned to the light blue-shaded time points is state 3. The cluster time courses (illustrated as color-shaded bars, over a single fMRI session) are then used to calculate the probability and the state switching probabilities of each PL state in each condition (e.g. music, no music). *Middle:* paradigm used in our study. *Bottom:* Given the cluster time courses, the probability of occurrence of each PL state and the switching probabilities in the different conditions (music vs no music) are calculated.

### fMRI image acquisition

All preadolescents underwent anatomical, diffusion weighted, and functional MR imaging of their brain. Data collection parameters only for the structural and functional scans are described here.

A 3 Tesla scanner (Siemens Magnetom Tim Trio), equipped with a 32-channel head coil, was used to obtain the images. To prevent postural adjustments and to attenuate the noise and vibration of the scanner, foam cushions were placed around the arms of the participants. Music was presented through audio headphones with active noise control (OptoACTIVE, Optoacoustic).

BOLD-weighted fMRI data were acquired using a gradient echo-planar imaging (EPI) sequence. The scans comprised 469 volumes of 48 axial-slices (TR: 1300 ms; TE: 27.32 ms; FOV: 192 x 192; voxel size = 2.50 x 2.50 x 2.50 mm3). During the fMRI measurement, participants listened to the stimulus presented at an average sound level of 80 dB. The participants were instructed to stay still and to relax during the whole fMRI paradigm without falling asleep. Subsequent to a short break after fMRI recording, a high-resolution T1-weighted 3D structural image was acquired for anatomical reference (176 slices; TR: 2420 ms; TE: 3.7 ms; FOV: 256 x 256; voxel size: 1 x 1 x 1 mm3). During the whole scanning session an MRI-compatible eyetracker (EyeLink 1000 – SR research) was used to better monitor the preadolescents and to ensure that they did not fall asleep during the fMRI paradigm.

### Preprocessing

The initial preprocessing of the fMRI data was performed with FSL (FMRIB’s Software Library, www.fmrib.ox.ac.uk/fsl) using the MELODIC tool (version 3.14). The following standard steps were performed within MELODIC:

1. Removal of the first five EPI volumes to allow for signal stabilization.
2. High-pass filtering >0.01Hz, removing signal components with periods longer than 100 seconds.
3. Spatial smoothing of the EPI volumes with an FWHM of 4 mm (Smith and Brady 1997).
4. Motion correction with MCFLIRT (Jenkinson et al. 2002).
5. Registration of the EPI data to standard MNI space through a dual linear registration process using FLIRT (Jenkinson and Smith 2001; Jenkinson et al. 2002), including a transformation between the EPI data and a structural T1 scan of the same participant and a transformation between the same structural T1 scan and an average brain template in MNI space for children ranging in age between 7.5 and 13.5 years (Fonov et al. 2011). Prior to the registration process participants’ T1 brain images were extracted using FSL’s brain extraction tool (BET) (Smith 2002).
6. Noise removal using probabilistic spatial independent component analysis (ICA) decomposing each participant’s fMRI session into N_components_ × time, with the number of components, N_components_, being estimated automatically through Bayesian dimensionality estimation techniques (Beckmann and Smith 2004) and each component being represented by a spatial map and a timecourse. The resulting ICA decompositions (one per participant) were reviewed independently by two raters (experimenters M.C.F. and A.S.) and classified into noise or signal based on criteria outlined in Griffanti et al. (2017) for the frequency content and appearance of the timecourse and for the pattern of the spatial map of each component. Following this hand classification, the FSL function *fsl_regfilt* was used to recompose the data to N_voxels_ × time while regressing out the contribution of the components labeled as noise.

After this pre-processing, we extracted the time courses of BOLD activity in 90 cortical and subcortical regions-of-interest (ROIs) defined using the Automated Anatomical Labeling (AAL) atlas (Tzourio-Mazoyer et al. 2002). This was done by registering the AAL template to the standard Fonov brain relative to which a transformation matrix existed to each participant’s native EPI space using FLIRT (see above). Representative timecourses were estimated using the FSL function *fsl_meants* as the mean across the voxels included in each ROI. The BOLD signals in each of the 90 brain areas were subsequently band-pass filtered between 0.02 and 0.1Hz (using a 2^nd^ order Butterworth filter as in Lord et al., 2019), discarding in this way the high frequency components associated to cardiac and respiratory signals (>0.1Hz), and focusing on the most meaningful frequency range for detecting functional connectivity in BOLD signal fluctuations (Biswal, Yetkin, Haughton, & Hide, 1995; J. Cabral et al., 2017; Glerean, Salmi, Lahnakoski, Jaaskelainen, & Sams, 2012).

### Dynamic Phase Locking Analysis

To evaluate the temporal dynamics of recurrent functional networks, we used BOLD Phase synchronization measures (Glerean et al. 2012; Ponce-Alvarez et al. 2015; Deco and Kringelbach 2016; Cabral, Vidaurre, et al. 2017; Deco et al. 2017). We first estimated the phase of the BOLD signals in all *N*=90 areas over time, *θ(n,t)*, using the Hilbert transform, which expresses a signal *X* as *X(t)=A(t)e^iθ(t)^*, where *A(t)* is the instantaneous amplitude, and *θ(t)* is the instantaneous phase. In Figure 1A we represent all *N=90* BOLD phases at time *t* in the unit circle in the complex plane, where the real axis corresponds to *cos(θ(n,t))* and the imaginary axis corresponds to *sin(θ(n,t)*).

To obtain the pattern of BOLD phase locking between all brain areas at each single time point *t*, we compute a dynamic Phase Locking matrix *dPL(n,p,t)*, with size *NxNxT*, where *N*=90 is the number of brain areas considered in the current parcellation scheme (see Preprocessing), and *T=464* is the number of recorded frames in each scan. The phase locking matrix between brain areas *n* and *p* at time *t* is obtained using the following equation:

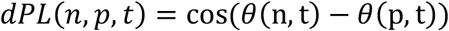

where *cos()* is the cosine function. *dPL(n,p,t)* is 1 if two areas *n* and *p* have synchronized BOLD signals at time t, and *dPL(n,p,t)* is 0 if the BOLD signals are orthogonal (with a phase difference of 90°).

### Leading Eigenvector of BOLD Phase-Locking

To characterize the evolution of the dPL matrix over time, we employed the Leading Eigenvector Dynamics Analysis (LEiDA) method which captures the temporal evolution of phase-locking patterns with reduced dimensionality (Cabral et al., 2017b). The leading eigenvector of the dPL matrix at time *t*, *V_1_(t)*, is a Nx1 vector that captures the main phase orientation over all areas, where each element in *V_1_(t)* represents the projection of the BOLD phase in each brain area into the leading eigenvector (Figure 1A) (Lord et al. 2019).

When all elements of the leading eigenvector, *V_1_(t)*, have the same sign, it means all BOLD phases are following the same direction with respect to the orientation determined by *V_1_(t)*, which is indicative of a global mode governing all BOLD signals (Figueroa et al. 2019; Vohryzek et al., 2020; Lord et al., 2019). If instead the first eigenvector *V_1_(t)* has elements of different signs (i.e. positive and negative), the BOLD signals follow different directions with respect to the leading eigenvector, which we use to divide brain areas into 2 ‘communities’ according to their BOLD phase relationship (see Figure 1A) (Figueroa et al. 2019). The magnitude of each element in *V_1_(t)* indicates the ‘strength’ with which brain areas belong to the communities in which they are placed (Newman, 2006).

Eigenvectors can be represented in cortical space by representing each element as a sphere placed at the center of gravity of the corresponding brain area, and scaling the color of each sphere according to its relative projection onto *V_1_(t)*: yellow-to-red for positive projections and cyan-to-blue for negative projections where lighter colors (cyan/yellow) indicate stronger contributions and darker colors (blue/red) weaker contributions (Figueroa et al. 2019). To highlight the network formed by the smallest community of brain areas (which we find to reveal meaningful functional networks, as shown in Vohryzek et al., 2020; Lord et al., 2019; Laraibi et al., 2020), we plot links between the corresponding areas. For example, Figure 1C shows five PL states represented in cortical space, where the BOLD signals can be divided into 2 modes according to their relative phase alignment: a larger set of brain areas (cyan areas) and a smaller functional network (orange/red areas) formed by areas whose BOLD signal is coherent but phase-shifted with respect to the larger community.

### BOLD PL states - Detection of recurrent patterns of whole-brain BOLD phase locking

Our aim was to explore whether there are specific PL configurations that differentiate between music listening and no music. To do so, we first clustered the whole sample of PL patterns into a varying number of recurrent patterns, applying a *k*-means clustering to all leading eigenvectors *V_1_(t)* across all preadolescents (7854 leading eigenvectors corresponding to all 462 TRs of all 17 preadolescents) (see Figure 1B). The clustering divides the sample into a *k* number of clusters (each representing a recurrent PL configuration), with higher *k* revealing rarer and finer-grained network configurations. The centroids of each cluster *c* are vectors with size Nx1, which we use to represent recurrent PL states. Importantly, the clustering assigns a single PL state to each fMRI time frame, as highlighted by the shaded bars in Figure 1D.

Since the exact number of functional networks remains under debate, we ran the *k*-means clustering algorithm with *k* ranging from 3 to 15 (i.e. dividing the sample of eigenvectors into k=3, k=4, …, k=15 clusters) to cover the range of functional networks commonly reported in literature (Damoiseaux et al. 2006; Yeo et al. 2011). Subsequently, following the approach of Figueroa et al. (2019), we examined, for each *k*, how each PL state changed between conditions, in order to detect any PL state that significantly differs in probability between music listening and no music.

### Occurrence and switching profiles of PL states

Upon identifying BOLD PL states, we computed the probability of occurrence of each PL state in each condition (i.e. music and no music). The probability of occurrence (or fractional occupancy) is simply the number of epochs assigned to a given cluster divided by the total number of epochs (TRs) in each experimental condition. The probabilities were calculated for each subject, in each experimental condition and for the whole range of clustering solutions explored as in Figueroa et al. (2019). In addition, we computed the switching matrix, which captures the trajectories of PL dynamics in a directional manner, namely indicating the probability of transitioning from a given PL state to another (Lord et al. 2019; Figueroa et al. 2019).

Differences in probabilities of occurrence and in probabilities of transition for the different states between music and no music were statistically assessed using a permutation-based paired t-test (10000 permutations). For each of the 10000 permutations, a t-test was applied to compare conditions.

To evaluate the significance of results taking into account the family-wise error rate (i.e. probability of false positives arising from multiple comparisons), we define two corrected thresholds of *α_1_*=0.05/*k* (green dashed line in Figure 2A/B) and *α_2_* = 0.05/ Σ(k)=0.05/117 (blue dashed line in Figure 2A/B). The first one, *α_1_*, takes into account the number of independent hypothesis tested in each partition model. The second threshold, *α_2_*, is the most conservative Bonferroni correction considering all hypothesis tested, but since the hypothesis are not independent across *k* (as we will show, some PL states are highly correlated across *k*), this threshold can be too conservative, increasing the probability of missing true positives (Figueroa et al, 2019).

**Figure 2.**
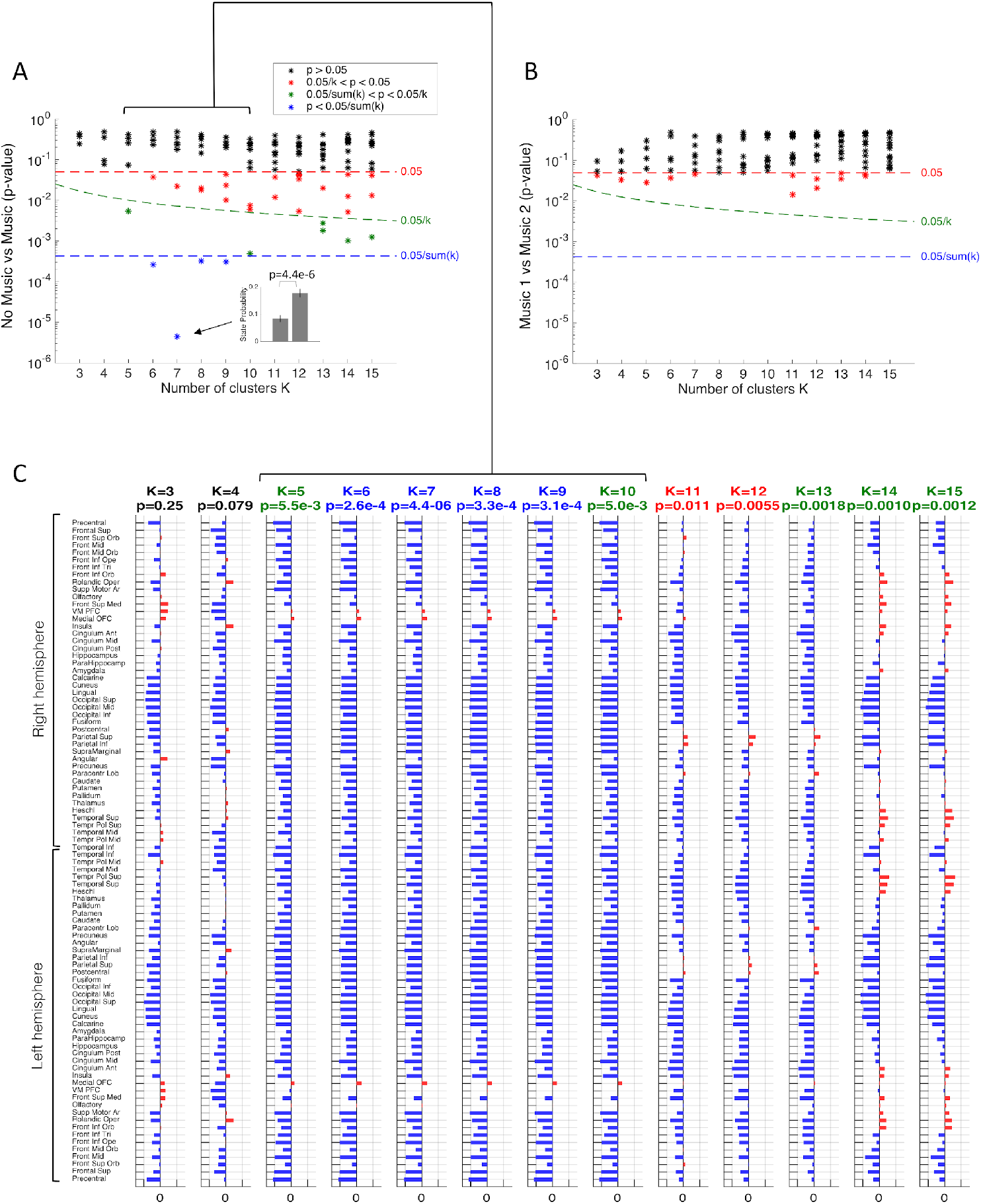
Probability of some PL states differs significantly between music and no music but not between similar music pieces, over the range of partition models explored. **A –** We compared the probability of occurrence of each PL state between Music and No music for each partition model (i.e. with number of clusters k varying between k=3 and k=15) calculating the corresponding *p*-values. For each *k*, we obtain *k p*-values (*), each corresponding to a different PL state. We colour each *p*-value according to its significance level, determined by its relative position with respect to the different threshold lines: the red dashed line represents the standard uncorrected threshold of 0.05, green is the threshold corrected by the number of hypothesis tested within each partition model as 0.05/*k*, and blue is the threshold corrected by 0.05/117, where 117 is the total number of comparisons performed. All *p*-values above 0.05 are coloured in black. We find a number of PL states that show highly significant differences between music and rest, some of which surviving the most conservative correction for multiple comparisons (blue dots). **B –** The same analysis was performed but this time comparing the probabilities of PL states between the two similar music pieces (Music 1 and Music 2) listened during the same scanning session. As can be seen, although some comparisons fall below the 0.05 threshold (red asterisks), these can be considered false positives due to multiple comparisons, meaning that the two medleys were well matched. **C –** For each partition model, we represent the PL state that most significantly differed between music and no music in vector format, with the corresponding *p*-value on the title. As can be seen, there is a robust PL pattern that consistently appears significantly different between conditions over a range of partition models (i.e. with *k*=5,6,7,8,9,10). In particular, for *k=7*, one of the 7 PL states shows a difference in probability between No music and Music with a *p*>-value=0.0000044 (< 5×10^−6^) (the PL state is shown in detail in Figure 4).

In order to verify the robustness of our results, we also measured the differences in probability of occurrence of PL states between piece 1 (Music 1) and piece 2 (Music 2).

As a further analysis, we correlated the overall BMRQ scores with the differences in occurrence and switching profiles of PL states between music and no music to better explore the neural underpinning of music reward sensitivity.

Moreover, we correlated the difference in probability of occurrence of BOLD PL states during music vs. no music with the preadolescents’ individual level of music training (determined in number of months of music education) and with the SES, with the purpose of understanding if previous music training experience and socio-economic status could affect the difference of PL configurations across the two conditions. The results of these additional analyses are presented in the *Supplementary material*.

## Results

### Detection of the most important BOLD phase-locking states

We first searched for BOLD PL configurations that significantly differentiate active music listening from no music. To do so, for each solution with *k* PL states (i.e. with number of clusters *k* varying between *k=3* and *k*=15), we compared the probability of occurrence of each PL state in each condition across subjects and obtained the corresponding *p*-values. In Figure 2A we show all the *p*-values obtained between conditions for each clustering solution (i.e. as a function of *k*). We found that, for most values of *k*, there is at least one PL state that significantly differs in probability between music and no music, passing the corrected threshold for the number of independent hypothesis (*<α_1_*, green asterisks) in 9 out of the 13 partition models (i.e. with *k=5,6,7,8,9,10,13,14,15*) and in 4 cases (*k*=6,7,8,9) surviving even the most conservative correction for positive-dependent multiple comparisons (*<α_2_*, blue asterisks).

The partition into 7 states revealed the most significant difference between music and no music, with one PL state showing increased probability of occurrence during music listening with *p* < 5×10^−6^. Importantly, as illustrated in Figure 2C, this result is not exclusive to the division into 7 states, with the partitions into *k*=5,6,7,8,9 and 10 states revealing a very similar PL pattern increasing its probability while listening to music. Indeed, the Pearson’s correlation between PL patterns obtained from *k*=5 to *k*=10 correlate with *r* = 0.98, which indicates that they refer to variant forms of the same underlying PL state, with differences arising from the number of output states constrained by *k*. Different PL states were found to vary significantly in probability when defining more fine-grained partitions into *k*=13, 14, 15, revealing more rare but also less statistically significant network configurations (>*α_2_*). Overall, the consistency of our findings for a range of partition models reinforces the existence of a specific pattern of BOLD phase locking that differentiates the effects of music listening from no music.

All the four partition models passing the most conservative Bonferroni correction returned a repertoire of PL states where only one PL state significantly differed in terms of probability between music and no music (see Figure 2A, blue dots), whereas all other PL states did not show differences surviving the correction for the number of independent hypothesis tested (p<0.05/*k*, see Figure 2A, red and black asterisks). For the subsequent analysis, we selected the partition into *k*=7 PL states since it is the one that detects the PL state that most significantly differs between music and no music in terms of statistical power (p < 10^−5^). Notably, the partition into 7 states is aligned with previous studies in the resting-state literature (Damoiseaux et al. 2006).

As a further validation analysis, we compared the probabilities of each PL state during the two very similar pieces of music that the preadolescents listened to during the scan (Figure 2B). In this case we did not detect any significant difference in probability of PL states between piece 1 and piece 2 over the whole range of partition models (all p<0.05/*k*, for all k between *k*=3 and *k*=15, red/black asterisks).

### Relevant PL state

Our analysis revealed a PL state, which consistently appeared more often during music listening than during the no music period (Figure 3 and PLstate.mov in *Supplementary Material*). This PL state consists of a network including bilateral medial orbitofrontal cortex (OFC), bilateral ventromedial prefrontal cortex, and left olfactory cortex. We further refer to this PL state as the ‘reward PL state’ since it involves key brain areas of the reward system. For the selected clustering model (*k*=7), the reward PL state occurred significantly more frequently in music than in the no music period (18±1.5% compared to 8±1.3% of the time, *p*=4×10^−6^ uncorrected, *p*=5×10^−4^ corrected by the 117 nonindependent hypothesis tested).

**Figure 3.**
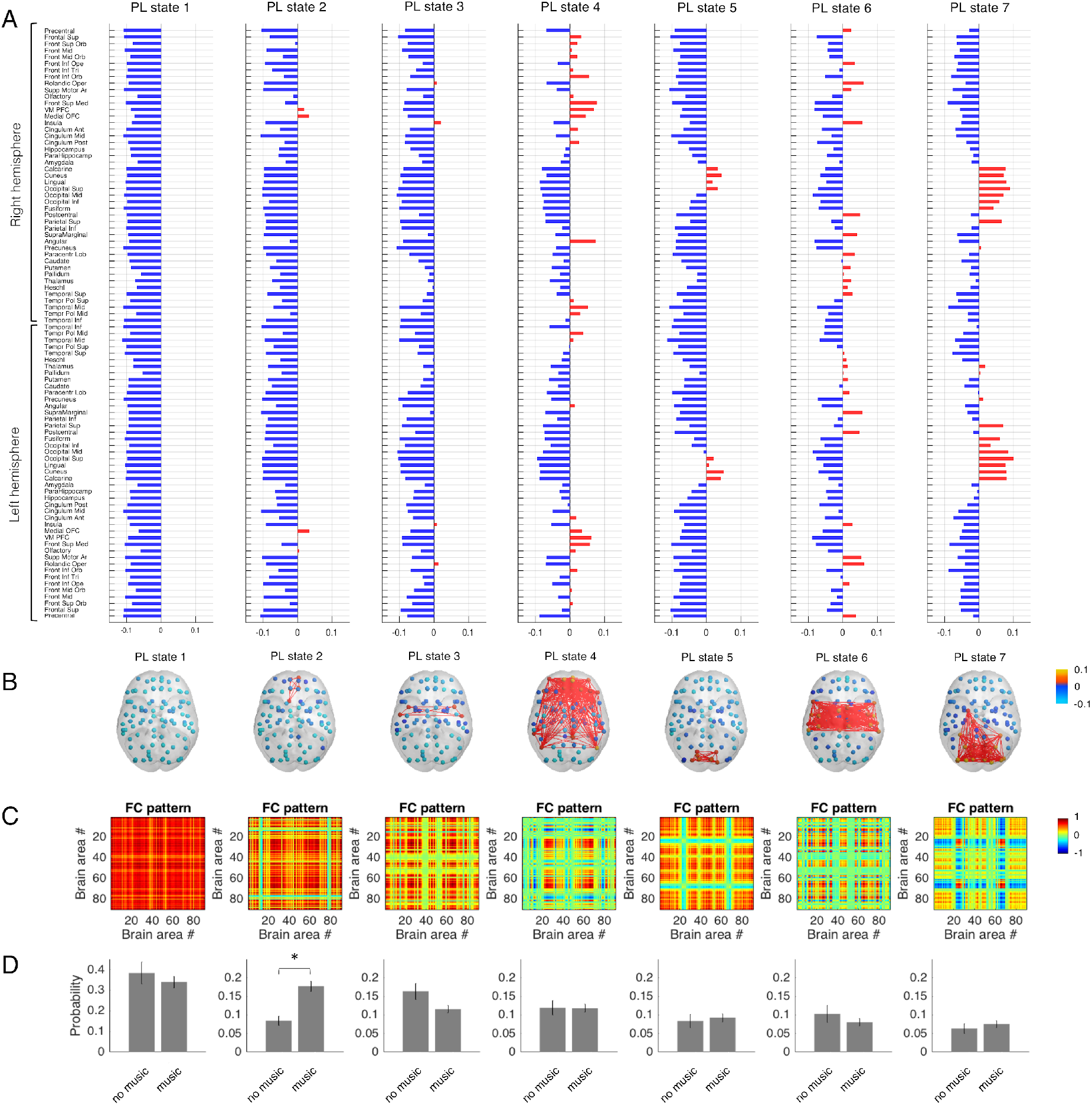
Repertoire of PL states assessed with LEiDA (for *k*=7) and their probabilities of occurrence during music compared to no music. PL states are represented as vectors (**A**), in cortical space (**B**) and in matrix format (**C**). **A –** Each PL state is represented by a central vector, *V_c_* (centroid) of each cluster of leading eigenvectors. Bar charts represent projections of each of the *N=90* AAL brain areas to each of the *k=7* PL states (*V*_c_*(n)>0* in red, blue otherwise; where *n=1, …, N* and *c*=1, …, *k*). **B –** Each *V_c_* is represented in cortical space by plotting spheres at the center of gravity of each brain area *n* and coloring them according to the value in *V*_c_(n): lighter and darker colors show stronger and weaker coherence, respectively, within the positive (yellow-red) vs the negative (cyan-blue) communities. **C –** Each *NxN* matrix (given by the outer product of each cluster centroid vector *V_c_V_c_^T^*) represents a recurrent dominant phase-locking pattern. **D –**Probability of occurrence of each PL state in each experimental condition (music and no music), with asterisks (*) denoting statistical significance with *p* < 0.05/k in the probability of occurrence between conditions. Only one PL state (2) shows significant difference in probability between music and no music (*p*=4×10^−6^ uncorrected, *p* = 5×10^−4^ Bonferroni corrected by the 117 non-independent hypotheses tested).

**Figure 4.**
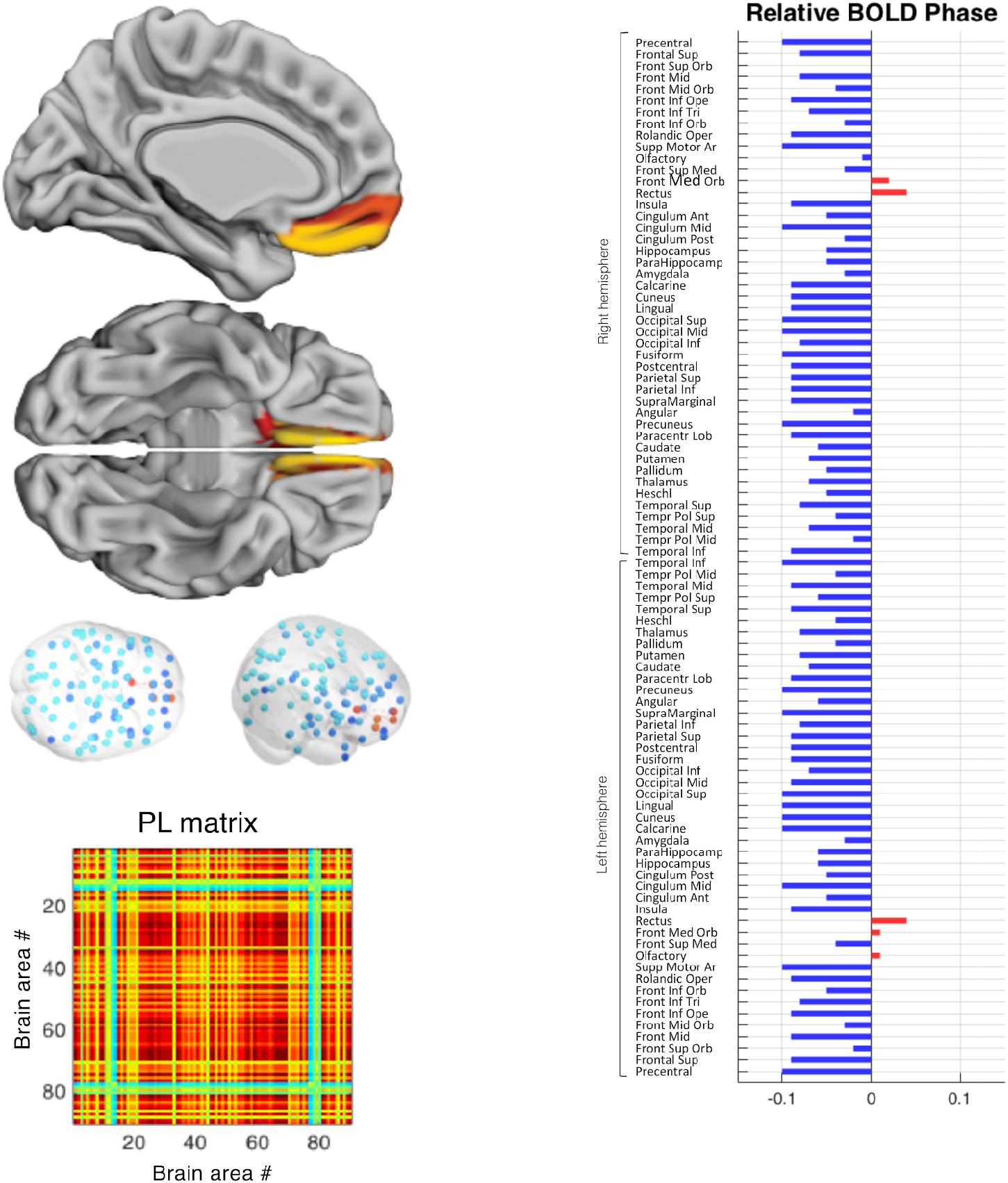
Characterization of the PL state occurring significantly more often during music listening than no music. The PL state for which fractional occupancy is increased significantly during music listening compared to no music corresponds to a network involving OFC bilaterally. *Left top and middle:* graphical representation of the functional network where links represent all positive functional connections between the most relevant nodes belonging to the reward network. *Left bottom:* PL pattern represented in matrix format as the outer product of the relevant cluster centroid vector *V_c_V_c_^T^. Right:* bar chart representing contributions of each of the *N=90* AAL brain areas to the PL state of interest (*V_c_(n)>0* in red, blue otherwise).

Figure 3 illustrates the full repertoire of PL states that are returned by LEiDA when choosing *k=7*. This reveals different network configurations that appear, dissolve and reoccur in all participants during the scan. In Figure 3D, we show the mean probability of occurrence (and standard error of the mean, SEM) of the 7 PL states in each condition. Except from PL state 2 (the ‘reward’ PL state), none of the other 6 PL states showed significant differences between music and no music. In alignment with previous studies in adults, a state of global BOLD phase coherence (PL state 1) is found to occur with the highest probability (Cabral, Vidaurre, et al. 2017; Lord and Roseman 2018). Notably, some of these networks show spatial overlap with previously-described resting-state networks (Yeo et al. 2011) and with cognitive states based on a large-scale automatic synthesis of human functional neuroimaging data (Yarkoni et al. 2011).

### Switching probabilities

We examined the transition patterns between PL states in detail for the selected partition model (*k*=7) by calculating the probability of, being in a given PL state, transitioning to any of the other states. In Figure 5 we illustrate the general switching pattern during no music (Figure 5A right) and music (Figure 5A left) and the difference between the two conditions (Figure 5 A/B). An illustration of music vs. no music changes in the transition probabilities between PL states rendered on the cortical surface is provided to facilitate the neuroanatomical interpretation of PL state switching (Figure 5 B), with red arrows indicating the switches that occurred more often in the music condition than in no music, and in blue the switches that were more frequent during no music.

**Figure 5.**
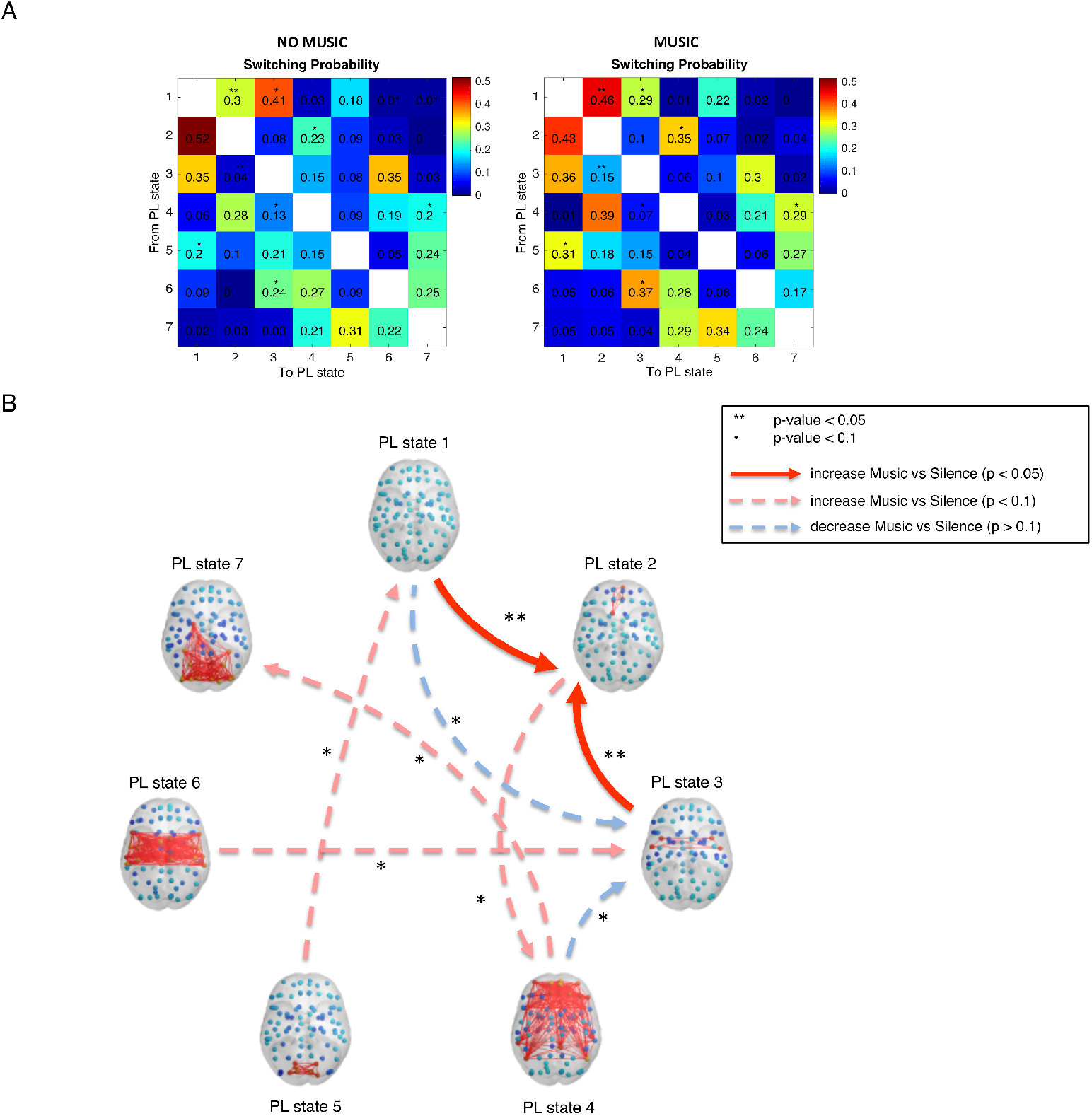
Switching probabilities during music listening and no music and differences between the two conditions. **A –** Switching matrices showing the probability of, being in a given PL state (lines), transitioning to any of the other states (columns) both during no music (left) and music (right). Significant between-condition differences assessed via a permutation test are indicated by two asterisks (**) for a significance threshold *p*<0.05 and with one asterisk (*) for a threshold *p*<0.1 **B –** Graphical illustration of the most significant differences between Music and No music in the transition probabilities shown in A. Each arrow represents a state-to-state increased/decreased transition probability in music compared to transition probability in no music; red arrows represent an increased probability of transition in music compared to no music, while blue arrows show reduced transition probabilities between states in music compared to no music.

We found that the probability of switching from PL state 3, involving bilateral insula and rolandic operculum, towards the reward PL state (PL state 2) was increased during music compared to no music (15% vs. 4%, *p*=0.006 uncorrected, *p*=0.04 corrected by *k*). Moreover, the state of global coherence (PL state 1) showed a statistical trend of transitioning more often to the reward PL state during music than during no music (46% vs. 30%, *p*=0.01 uncorrected).

### Correlation with individual musical reward sensitivity

We correlated the *difference* in probability of occurrence of the reward PL state between music and no music for *k=7* with the music reward sensitivity of the preadolescents, as assessed by means of the BMRQ questionnaire, and we found a non-significant correlation (*p*>0.05).

We subsequently examined the relationship between the switching probabilities between PL states for the selected partition model (*k*=7) and the individual BMRQ scores. In Figure 6 (*left*), the matrix values refer to the correlation between the individual difference of switching patterns between music and no music and the individual BMRQ score. Only one transition showed a correlation with the BMRQ score with *p*<0.05 referring to an increased probability of switching from PL state 3 towards the reward PL state (PL state 2) during music compared to no music. The scatter plot of the individual switching probabilities (x axis) versus the individual BMRQ score (y axis) is reported (Figure 6, *middle*), together with an illustration of the transition (red arrow) from PL state 3 to PL state 2 rendered on the cortical surface to facilitate the neuroanatomical interpretation of the relationship between BMRQ and PL state switching (Figure 6, *right*).

**Figure 6.**
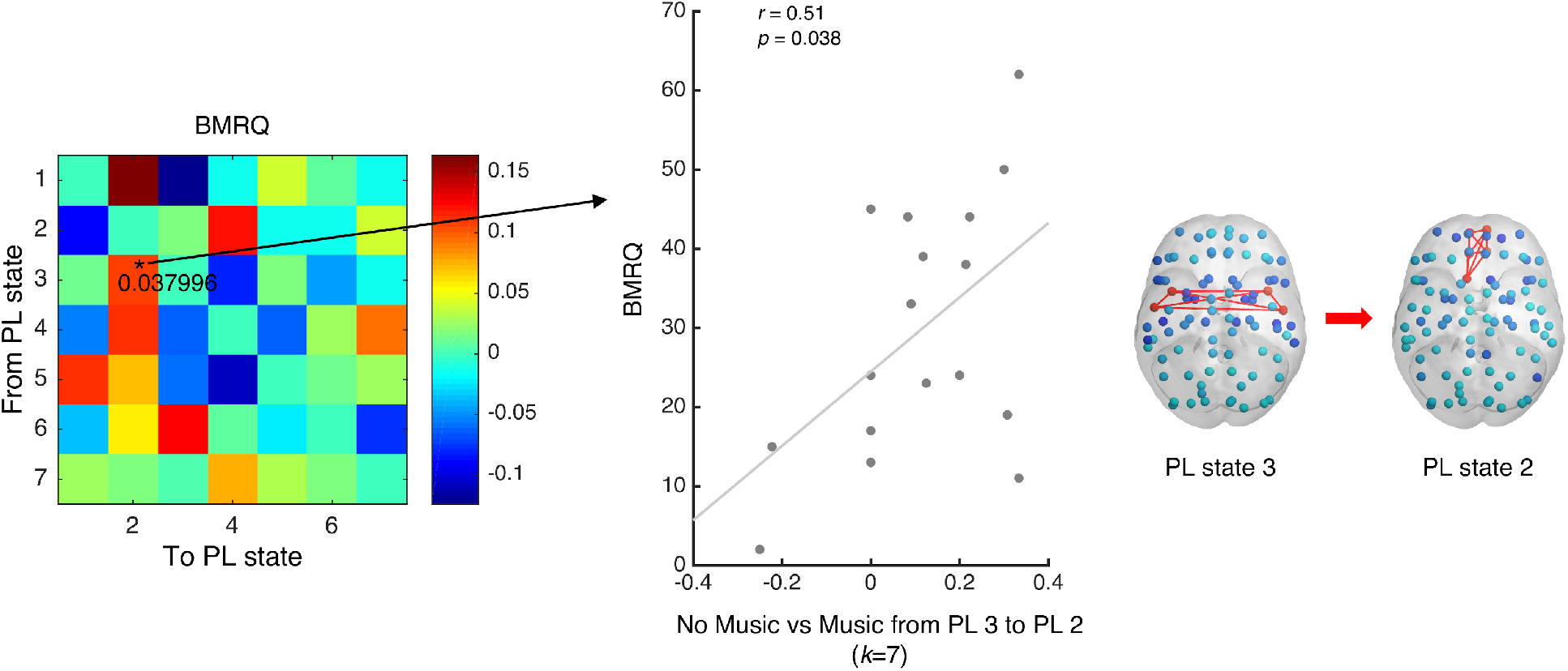
Correlations between musical reward sensitivity (BMRQ) and (A) differences in reward PL state probability or (B) differences in switching probabilities between music listening and no music. *Left:* Switching matrix showing the correlation between BMRQ score and the difference between music and no music in probability of, being in a given PL state (lines), transitioning to any of the other states (columns). Significant between-condition difference assessed via a permutation test is indicated by one asterisk (*) for a significance threshold *p*<0.05. *Middle:* Scatter plot of individual BMRQ scores and differences between Music and No music in the transition probabilities from PL states 3 to 2. *Right:* Graphical illustration of the state-to-state transition (red arrow) occurring with increased probability during music in participants with high BMRQ score than with low BMRQ score.

## Discussion

We here identified a significantly increased occurrence of a BOLD Phase-Locking state during music listening compared with no music in preadolescents (10-11 years of age), consisting of a pattern where the BOLD signals of bilateral medial OFC and ventromedial prefrontal cortex are phase shifted from the rest of the brain. The OFC involved in this PL state is considered to be a key area of the brain’s pleasure system and has been consistently related to musical reward (Blood et al. 1999; Blood and Zatorre 2001; Trost et al. 2012; Salimpoor and Zatorre 2013). Similarly to other pleasant activities, music listening has been associated with a cyclical time course of pleasure including: a phase of expectation or wanting for a specific rewarding musical structure; a phase of consummation or liking of the music reward, which can have a peak level of pleasure (e.g., musical chills); a satiety or learning phase, where one learns and updates musical predictions changing both the wanting phase and the liking phase for future listening experiences (Gebauer et al. 2012; Georgiadis and Kringelbach 2012; Kringelbach et al. 2012; Brattico et al. 2013; Brattico 2015, 2019a). These musical pleasure cycles, driven by different mechanisms including musical expectancy, memory associations, evaluative conditions (for a review, see Gebauer et al. 2012; Brattico 2019b), involve the reward brain system, in particular the OFC, the ventral tegmental area, and the nucleus accumbens (Blood and Zatorre 2001; Brown et al. 2004; Menon and Levitin 2005; Koelsch et al. 2006; Suzuki et al. 2008; Osuch et al. 2009; Brattico et al. 2015; Liu et al. 2016, 2017; Reybrouck et al., 2018). All these areas are similarly engaged in other pleasurable experiences involving food (Kringelbach et al., 2012) or sex (Georgiadis & Kringelbach, 2012). Although findings from MEG have indicated that the OFC plays a significant role on many timescales within the pleasure cycle (Kringelbach et al. 2008; Parsons et al. 2013; Young et al. 2016), this region has been proposed to be part of the top-down neural route to conscious liking of music, providing a modulation of the sensory musical pleasure by means of reward value attribution, as complementary to the bottom-up neural route to sensory pleasure in music operating in a fast manner without conscious awareness (Brattico 2015).

Despite the well-documented impact of music on the brain’s pleasure system and the important role that music listening plays in the lives of children and adolescents as a source of enjoyment (Giacometti et al. 1981; North et al. 2000; Erkkila and Saarikallio 2007; Miranda and Claes 2009), the neuroimaging studies focused on music listening conducted thus far have been focused mainly on adults. The only study with music listening in children that has been done by now has showed, by means of a general linear model, increased BOLD signal in the OFC in 10-year-old children when exposed to irregular chords (unexpected harmonic functions) vs. regular chords (Koelsch et al. 2005), corroborating the hypothesized ability of musical expectancy violation to engage emotional processing (Kraehenbuehl and Meyer 1957; Krumhansl 1997). However, the stimuli used in that study consisted of artificial chord sequences, therefore partially missing ecological validity. Our study is the first using a *free listening* task in preadolescents allowing them to listen and to enjoy real music (rather than very brief or artificial sounds) without being interrupted by unnatural behavioral tasks (Brattico 2019b; Alluri et al. 2012; Poikonen et al. 2016; Haumann et al. 2018).

A second aspect of novelty of this study relies on the employment of a new method of analysis able to measure the dynamic processing of music listening. The temporally extended musical pleasure cycle during music listening (Koelsch et al. 2006; Gebauer et al. 2012; Brattico et al. 2013; Brattico 2019a), underlines the need of understanding the neural underpinning of music listening using a method that allows to catch the dynamic networks that re-occur and dissolve over an extended period of time. Using LEiDA, we were able to explore music processing focusing on time-varying wholebrain BOLD connectivity patterns rather than relying on static correlational patterns or seed-based approaches.

The ecologically valid paradigm, in combination with the LEiDA methodology adopted to measure the dynamic processing of music, allowed us to capture the potential neural base hidden behind the well-established tendency of children and adolescents to spend hours listening to music (Tarrant et al. 2000; Erkkila and Saarikallio 2007; Roberts et al. 2009).

We show that music is able to attract recurrently in preadolescents a reward brain network involving a region that is strongly involved not only in musical pleasure cycles but in the general experience of reward. Our additional analysis comparing the probabilities of PL states between the two similar music pieces listened during the scanning session did not reveal any significant difference, supporting the replicability of our results. After the *free listening* paradigm, the participants indicated an average liking of 3.5 on a 5-point Likert scale of how much they liked the pieces they had listened to, showing to perceive the pieces as overall pleasurable. However, further studies may use the favorite pieces of each participant as stimuli, in order to further explore the attractive power of music to recruit brain reward systems when listening to familiar and highly rewarding stimuli.

Additionally, we observed a higher probability to switch from both the PL state 1 (global mode) and the PL state 3 (insula and rolandic opercuum) to the reward PL state during music listening compared with no music. Interestingly, we found a positive correlation between the music reward sensitivity of the participants, measured by means of BMRQ, and the tendency to switch from PL state 3 to the reward network (PL state 2): the preadolescents more inclined to be rewarded by music switched more frequently from the state involving insula and rolandic operculum, to the state involving the medial OFC reward system. Similarly to our results, Koelsch and colleagues (Koelsch et al. 2005) found that the rolandic operculum was recruited in conjunction with the anterior superior insula while children were listening to pleasant music and the authors related the activity in this network, together with the ventral striatum, to the emotional perception of the musical stimuli. Considering the strong involvement of insula and OFC in the pleasure system, it is likely that the bilateral insula played a role in recruiting the activity in bilateral rolandic operculum during emotional perception of the musical stimuli before switching to a PL state involving OFC, especially in the most rewarded-by-music participants. The relationship between this transition and the music reward sensitivity is consistent with a previous study in which greater activity in medial OFC while listening to favorite music was correlated with self-reported capacity to experience pleasure in a range of different situations (Osuch et al. 2009).

The recruitment of the brain network involving OFC during listening to pleasant music can potentially provide new knowledge in the field of child and adolescent psychiatry. This complex brain region is intimately involved in all parts of the pleasure cycle and plays a crucial role in emotion regulation and cognitive control (Kringelbach and Rapuano 2016). In line with this, the immaturity of the OFC in adolescence has been related to the increased emotional reactivity that this period is characterized by (Monk et al. 2003; Thomas et al. 2004; Casey et al. 2008). OFC is one of the last brain regions to mature (Gogtay et al. 2004; Kringelbach 2005; Shaw et al. 2008; Tamnes et al. 2010) and its protracted development, in terms of synaptic pruning and myelination, suggest that it might not be able to provide sufficient top-down control of robustly activated reward and affect processing emotional regions (e.g., nucleus accumbens and amygdala) during adolescence (Galvan et al. 2006; Casey et al. 2008; Steinberg 2010a, 2010b; Strang et al. 2013). This developmental pattern increases the probability of risky decision making and high emotional reactivity, especially in those adolescents prone to emotional reactivity (Casey et al. 2008). In particular, abnormalities of the volume, the activity and the connections of OFC with subcortical regions strongly related to emotions such as amygdala, have been shown to be predictive of obesity, internet gambling disorder, cannabis or alcohol abuse in adolescents (Chai et al. 2011; Maayan et al. 2011; Peters et al. 2015; Cheetham et al. 2017).

Therefore, considering the crucial role of OFC in adolescence, our results might offer important insights at different levels. First of all, the remarkable ability of music to recurrently engage a brain state involving the medial OFC, known to be involved in liking, monitoring, learning and memory of the reward value of reinforcers, confirms the involvement of this brain region in the musical pleasure cycles, investigated through the lens of dynamic functional connectivity, in preadolescents. Moreover, the top-down control function of the OFC in the emotional network, although not yet fully matured in adolescence, and the frequent recruitment of a PL state involving this region during music listening, potentially provides the basis for using music listening as a coping strategy for emotion regulation in adolescence (Giacometti et al. 1981; Arnett 1995; North et al. 2000; Schwartz and Fouts 2003; Miranda and Claes 2009). We may speculate that the capability of music listening to stimulate and strongly attract the reward OFC reward system during adolescence might make music able to modulate the development and the function of top-down reward regulation of this brain region. This would be consistent with studies showing the effects of music training - and therefore continuous music exposure - on cortical thickness maturation in OFC (Hudziak et al. 2014) and inhibitory control (Moreno et al. 2011; Alemán et al. 2017; Holochwost et al. 2017; Fasano et al. 2019) in children and adolescents. Considering the vulnerability of this transition phase of life, the impact of music listening on the reward brain network should be considered in the clinical context. Indeed, since orbitofrontal dysregulation appears to be a signature feature across a broader field of psychopathology in adolescence, the potential role of music listening in modulating the functioning of OFC at this age might have potential implications for future therapeutic and/or preventive music intervention.

## Supporting information

Supplementary analyses

PLstate2

## Acknowledgement

We gratefully acknowledge all the preadolescents participating in this study and their parents. We also thank other members of our team who helped with data collection and data entry, in particular Ida Siemens Lorenzen, Johanna Pardon, Louise Ahle Jensen, Filo Ullrich, Laura Risager Ubbesen. We wish to thank also the elementary schools for their collaboration, Line Gebauer for her support in the preparation of the ethics protocol, Bjørn Petersen for helping with the recruitment, and Niels Trusbak Haumann for his assistance with the preparation of the fMRI paradigm. The authors declare that there is no conflict of interest.

## Funding

This work was supported by the Center for Music in the Brain (MIB), funded by the Danish National Research Foundation (DNRF 117). MLK is supported by the ERC Consolidator Grant: CAREGIVING (n. 615539), MIB (DNRF 117), and Centre for Eudaimonia and Human Flourishing funded by the Pettit and Carlsberg Foundations. JC is supported by Portuguese Foundation for Science and Technology CEECIND/03325/2017, Portugal.

## Notes

### Competing Interest Statement

The authors have declared no competing interest.

