## Supplementary analyses for "The early adolescent brain on music: analysis of functional dynamics reveals engagement of orbitofrontal cortex reward system"

### Correlation with months of music training

Since in our sample we had preadolescents with different levels of music education, for all the partition models (i.e., with $k$ ranging from 3 to 15) we examined whether the differences in probability of occurrence of PL configurations between music and no music were correlated with the number of months of music training that each preadolescent had. As shown in Supplementary Figure S1, the correlation survived the correction for the number of independent hypothesis ($p<0.05/k$) only for $k=6,14,15$ for a PL state representing a global mode of BOLD activity (respectively $p=0.048$, $p=0.042$, $p=0.045$; corrected). In the preadolescents with more months of music training the probability of occurrence of this PL state was higher during music compared to no music. Since we did not find a significant correlation between music training and difference in probability of occurrence of the reward PL state in the two conditions, we can deduce that music training was not related to the higher probability of occurrence of the reward PL state (PL state 2 for $k=7$) during music compared to no music.

A recent fMRI study using the same methods we used for our analysis (Cabral et al. 2017), found a clear relationship between the occurrence of the global mode state and cognitive performance, suggesting a potential role of global BOLD coherence in cognitive processing. Considering the well-documented ability of music training to enhance cognitive abilities in children and preadolescents (for a review see Hallam 2010), and the tendency observed in recruiting this state while listening to music, especially for the more trained participants, our results should be further investigated.

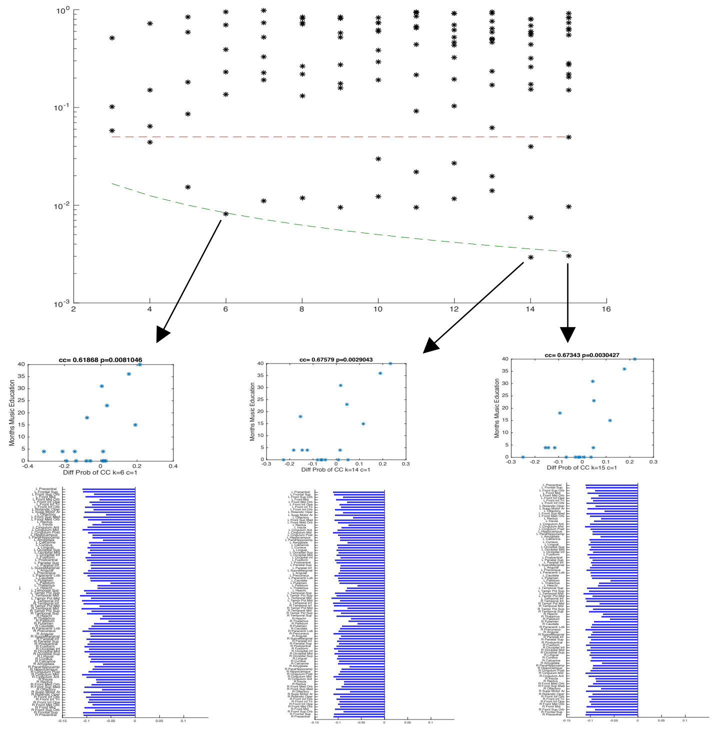


**Figure S1. Correlations between difference in PL state probabilities (between music and no music) and months of music training over the range of partition models explored.** The finding of a significant correlation between PL state 1 and month of music training was persistent over a range of partition models (*k*=6,14,15).

### Correlation with socio-economic status

We asked all the parents to indicate their highest level of education and annual household income on a questionnaire. For all the partition models (i.e., with *k* ranging from 3 to 15) we examined whether the differences in probability of occurrence of PL configurations between music and no music were correlated with the SES of the families of our participants. The results did not yield any significant correlation, meaning that the differences of pattern of PL-states elicited in music compared to no music were not linked to the SES.

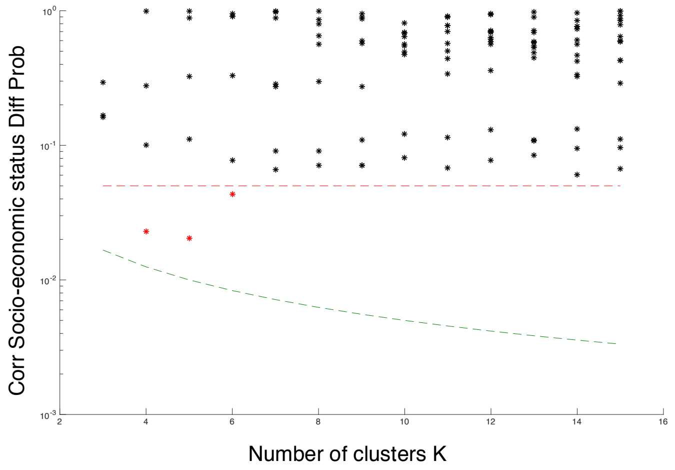


**Figure S2. Correlation between difference in PL state probabilities (between music and no music) and SES over the range of partition models explored.** No significant correlations was found.
